# Epigenetic regulation of differentially expressed genes between various glioma types

**DOI:** 10.1101/2020.08.29.272013

**Authors:** Ilona E. Grabowicz, Bartek Wilczyński, Bożena Kamińska, Adria-Jaume Roura, Bartosz Wojtaś, Michał J. Dąbrowski

## Abstract

Gliomas are the most frequent primary tumors of the central nervous system (CNS) and encompass two major subgroups: diffuse, malignant gliomas and benign, well differentiated gliomas showing a more circumscribed growth. Genome-wide next generation sequencing studies have uncovered specific genetic alterations, transcriptomic patterns and epigenetic profiles associated with different types of gliomas improving tumor diagnosis and having important implications for future clinical trials and patient management. We have recently created a unique resource encompassing genome-wide profiles of open chromatin, histone H3K27ac and H3Kme3 modifications, DNA methylation and transcriptomes of 33 glioma samples of different grades. Here, we took advantage of a wealth of data from those high-throughput experiments, intersected those data with topologically associating domains (TADs) and demonstrated that the chromatin organization and epigenetic landscape of enhancers have a strong impact on genes differentially expressed in low grade versus high grade gliomas. We identified TADs enriched in glioma grade-specific genes and/or epigenetic marks. We found a set of transcription factors, including REST, E2F1 and NFKB1, that are most likely to regulate gene expression in multiple TADs, containing glioma-related genes. Moreover, many genes associated with the cell-matrix adhesion Gene Ontology group, in particular 14 *PROTOCADHERINs*, were found to be regulated by the long range contacts with enhancers. Overall, the results presented here demonstrate the existence of epigenetic differences associated with chromatin organization driving differential gene expression in gliomas of different malignancy. We demonstrated that integration of whole genome epigenetic data with Hi-C data and transcriptomic profiles described in this work, can segregate low and high grade gliomas and reveal new regulatory networks that could explain some of the functional differences between gliomas of different malignancies.

**Highlights:**

- Integration of ATAC-seq, ChIP-seq and RNA-seq reveals glioma malignancy-related gene regulatory networks.
- TADs segmentation contributes to gene-epigenetically modified enhancer relationships.
- Contacts of active enhancers in gliomas of different malignancies might affect expression of genes involved in cancerogenesis, such as *PROTOCADHERINs* or *EGFR.*

## INTRODUCTION

Gliomas are tumors originating from neural stem cells or progenitor cells (Altmann et al. 2019). They range from benign and highly-curable pilocytic astrocytomas (World Health Organisation, WHO grade I, GI), through diffuse astrocytomas that could be benign (WHO grade II, GII) or malignant (WHO grade III, GIII) to highly malignant glioblastomas (WHO grade IV, GIV) (Uhlmann et al. 2003). Glioblastoma remains an incurable disease with a median survival of 15 months upon treatment and 3 months without treatment (Thakkar et al. 2014). Certain genetic alterations are currently used in glioma classification and survival prognostication, such as 1p/19q co-deletion, *ATRX* and *IDH1/2* mutations (Louis et al. 2016). *IDH* mutated gliomas display specific alterations of DNA methylation patterns (Turcan et al. 2012). Comprehensive studies of The Cancer Genome Atlas (TCGA) consortium have revealed that many epigenetics-related genes including chromatin modifiers, histone demethylases and deacetylases are being mutated in high grade gliomas (Brennan et al. 2013).

Epigenetic alterations occur in different types of cancers including gliomas (Burgess et al. 2008, Skiriutė et al. 2013, Ceccarelli et al. 2016, Dabrowski and Wojtas 2019, Barańska 2013). For example methylation of the *MGMT* gene promoter DNA in gliomas leads to gene silencing and is a favorable prognostic marker for patients treated with temozolomide (Burgess et al. 2008). Methylation of a set of specific cytosines allows for highly accurate prediction of overall survival of glioma patients from TCGA datasets (Dabrowski et al. 2018). Other epigenetic marks may activate or inhibit transcription. Acetylation of lysine 9 and 27 of histone 3 (H3K9ac and H3K27ac) and methylation of lysine 4 and 36 of histone 3 (H3K4me and H3K36me) are associated with active chromatin. Methylation of lysine 9 or 27 of histone 3 (H3K9me and H3K27me3) and DNA methylation in the genomic regulatory regions (such as promoters or enhancers) are the repressive marks (Burgess et al. 2008). The epigenetic marks regulate gene expression and control different transcriptional programs (Hirabayashi and Gotoh, 2010). The recent study of Stępniak et al. (2020) characterized the landscape of open chromatin and histone marks in gliomas of different grades providing a rich resource for further exploration. Gene expression patterns in glioma of different grades have been correlated with epigenetic marks depositions in the transcription start site (TSS) regions. In particular, the higher signals of H3K4me3 has been observed in pilocytic astrocytomas (PA) than in diffuse astrocytomas (DA) and glioblastomas (GB). In this study we took advantage of this data resource (Stępniak et al. 2020).

Chromatin openness, assayed with DNAse-seq or ATAC-seq assays, correlates positively with transcriptional activity (Dwivedi et al. 2019, Przanowski et al. 2019). Open chromatin - marked with H3K4me1 but depleted of H3K4me3, has been associated with enhancers (Heintzman et al. 2007, Smallwood and Ren 2013). Enhancers contain numerous transcription factor (TF) binding sites with their characteristic TF-motifs. TFs can act in either generic or cell-type specific manner (Buecker and Wysocka, 2012). Active enhancers are enriched for H3K27ac while repressed ones for H3K27me3 (Creyghton et al. 2010, Rada-Iglesias et al. 2011).

Another layer of gene regulation complexity is the organisation of the genome into three-dimensional domains, called topologically associating domains (TADs). TADs borders align with H3K27me3 or H3K9me2 blocks, lamin-associated domains as well as coordinately regulated gene clusters (Nora et al. 2012). They are stable across different cell types, highly conserved across species (Dixon et al. 2012) and their disruption can lead to aberrant contacts between genes and enhancers resulting in developmental diseases (Lupianez et al. 2015) and cancer (French et al. 2013, Johnston et al. 2019). Mutations in the CTCF motif within TAD borders, may affect its binding affinity and change gene expression of sets of genes (Umer et al. 2016). IDH mutation changing DNA methylation and/or histone methylation may lead to CTCF binding dysfunction in gliomas causing aberrant *PDGFRA* activation (Flavahan et al 2016).

Here, we take advantage of the wealth of data from high-throughput experiments: ATAC-seq, H3K4me3, H3K27ac ChIP-seqs, DNAse-seq, DNA bisulfite sequencing (providing information about DNA methylation) and RNA-seq acquired from glioma bulk tumour samples of various WHO grades (GI-GIV). We demonstrate that the epigenetic landscape of promoters and enhancers regulates expression of malignancy-specific genes. We found TADs unexpectedly enriched in glioma grade specific genes and/or epigenetic marks. Moreover, we identified a set of TFs which putatively regulate gene expression in multiple TADs, including known glioma related TFs such as REST, E2F1 and NFKB1. Many genes associated with the cell-cell adhesion Gene Ontology group, including *PROTOCADHERIN* family of genes, were found to be regulated by the long range contacts defined by the previously published Hi-C experiments performed on the human, developing brains (Won et al. 2016). Importantly, we found a large set of *PROTOCADHERIN* coding genes regulated by just one differentially acetylated enhancer.

## RESULTS

### Patterns of epigenetic marks depositions in differentially expressed genes between benign vs malignant gliomas

First, we determined differentially expressed genes (DEGs) between gliomas of different WHO grades GI-GIV. We compared: i) pilocytic astrocytomas (PA, WHO GI) vs diffuse astrocytomas (DA, WHO GII/GIII) (PA vs DA), ii) PA vs glioblastoma (GB, WHO GIV) and pediatric glioblastoma (PG, WHO GIV) samples (PA vs GB/PG) and iii) DA vs GB/PG. The numbers of DEGs were obtained for each of the three analyses: 2954 for PA vs DA, 4216 for PA vs GB/PG and 117 for DA vs GB/PG (DeSeq2, FDR-corrected p < 0.01). Some DEGs have also significantly differential epigenetic marks (DEMs) depositions at the promoter regions (DeSeq2, FDR-corrected p < 0.01). The majority of DEGs having deposited DEMs at the promoters were discovered for PA vs GB/PG comparison: there were 75 DEGs with H3K27ac DEMs and 87 with H3K4me3 DEMs (Figure 1A). The observed, significant intersection of DEMs with DEGs suggests an important role of the epigenetic regulation of genes involved in gliomagenesis when compared to the randomly selected, active genes (p<0.01, permutation testing). For PA vs DA we observed similar results (Figure S1A), in contrast to DA vs GB, where not many DEGs and genes with DEMs were found (Figure S1B).

**Figure 1.**
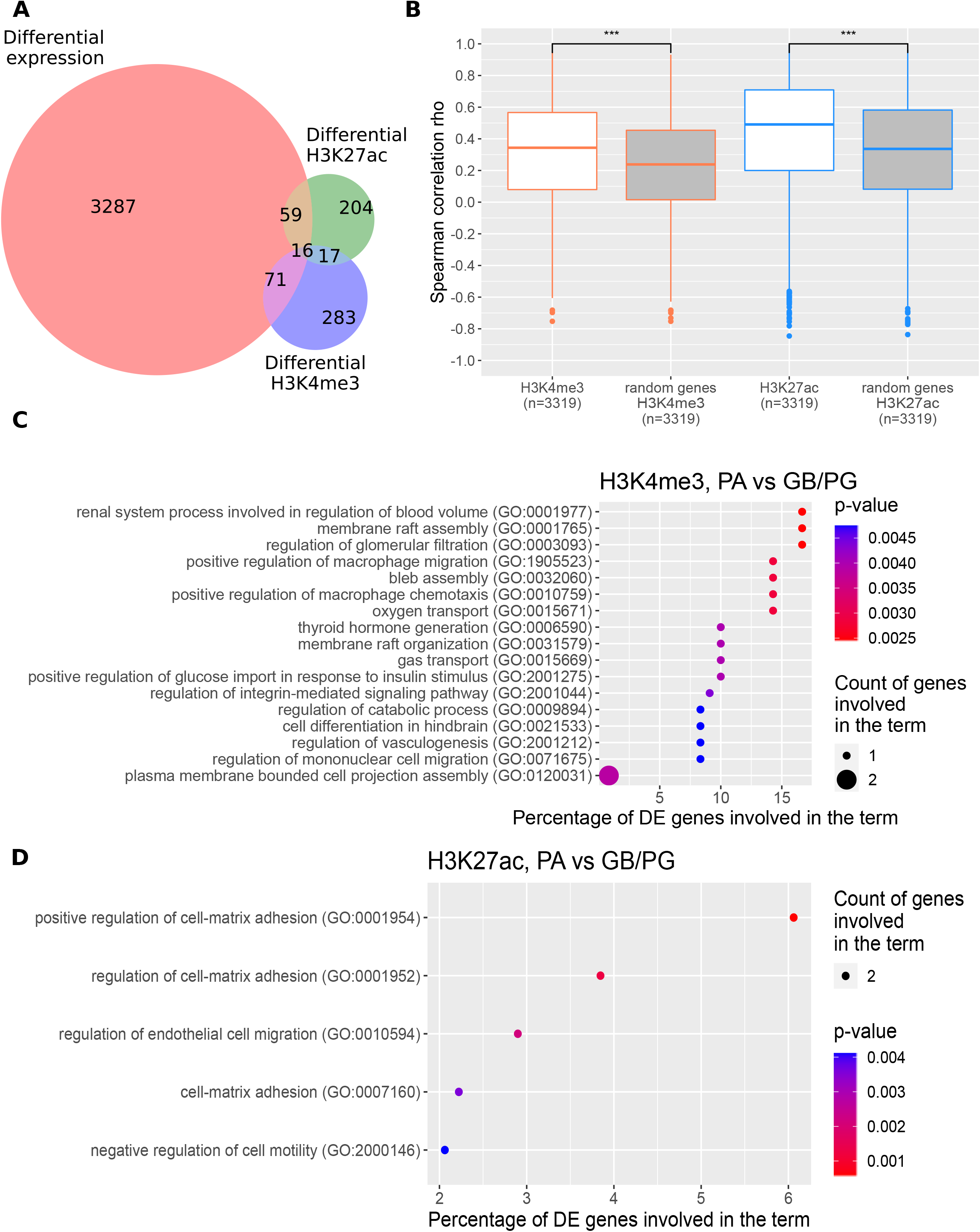
Identification of genes differentially expressed in benign and malignant gliomas (PA vs GB/PG). (A) Intersection of differentially expressed genes (DEGs) with genes carrying differential epigenetic modifications (DEMs) for the PA vs GB/PG comparison. (B) Correlation of H3K4me3 (orange boxes) and H3K27ac (blue boxes) coverages at the promoters of DEGs (PA vs GB/PG) with their expression. Boxes filled with white show values for DEGs, while in grey for the randomly chosen active genes. (C) Enrichment of Biological Process GO terms for DEGs in PA vs GB/PG comparison having high correlation of expression levels with H3K4me3 (Spearman rho > 0.7), and being prognostic for glioma patients’ survival (log-rank test p < 0.001). (D) As in Figure 1C for H3K27ac.

For further comparison of three glioma grades, DEGs and genes with assigned DEMs were selected. We found significantly stronger correlation between expression of DEGs and H3K4me3 and H3K27ac deposition at their promoters (0.31 - 0.42 and 0.57 - 0.67 for H3K4me3 and H3K27ac, respectively) than in the randomly selected non-DEG, active genes (Figure 1B; Figures S1C-S1D). The differences between DEGs/DEMs and randomly selected active genes reached statistical significance for all compared grades (Wilcoxon test, p < 9e-187). These results indicate a strong influence of epigenetic marks deposition in the gene promoters on transcription of genes involved in the development of glioma.

Next, we checked whether DEGs with the strongest correlation between gene expression and epigenetic mark levels are of clinical importance. When applying correlation threshold of Spearman rho > 0.7 we obtained sets of DEGs with expression levels highly correlated with epigenetic marks: 882, 41, 895 for H3K27ac and 491, 19, 383 for H3K4me3 for three pairwise comparisons: PA vs DA, DA vs GB/PG and PA vs GB/PG, respectively. The average share of the genes prognostic for patients survival (p<0.001 log-rank test, proteinatlas.org) among the highly correlated genes, for all three grade-differential-analyses was 1.28% (Table 1), while for all DEGs was just 0.34%. The observed difference reached a statistical significance (Wilcoxon test p < 0.05). This result shows that differences in epigenetic mark levels between tumors of various malignancy grades can affect the expression of key cancer genes and could make a difference in survival of glioma patients. Functional enrichments of those genes are connected to cancerogenesis (Figures 1C-1D, Figures S1E-S1H).

**Table 1.** DEGs with expression levels correlating with H3K4me3 or H3K27ac at the level of >0.7 Spearman rho and being highly prognostic for glioma patients survival (p<0.001 log-rank test, proteinatlas.org)

| Assay, comparison | DEGs, correlation$>0.7$, prognostic for patients survival |
| --- | --- |
| H3K4me3, PA vs DA | *SNX10, PHGDH, QPRT, CPQ, AEBP1, ADAMTSL4, UBXN10, EN2, CARD19, HTRA1, ISG20, GPC5, MKRN3, ACHE, MXRA5, KCNN4, RARRES2, LOXL2, MSTN, ETNK2, RNF175, FBXO27, CYGB, RPL39L, ETNPPL, FOSL1, HIST3H2A, CASC10, EMP2, FBXO17* |
| H3K4me3, PA vs GB/PG | *CPQ, UBXN10, MXRA5, RARRES2, SYT5, CYGB, CEND1, EMP2* |
| H3K4me3, DA vs GB/PG | *ISG20, GLUD1, PDGFA* |
| H3K27ac, PA vs DA | *MTHFD2, SNX10, PHGDH, QPRT, CPQ, AEBP1, LOXL1, DNAJA4, CTF1, CRELD1, EN2, CARD19, SMPD1, ISG20, MKRN3, ACHE, RARRES2, LOXL4, MSTN, ETNK2, RNF175, GLUD1, FBXO27, CYGB, ETNPPL, C11orf24, GRIK1, HIST3H2A, RP11-195F19.5, CASC10, FBXO17* |
| H3K27ac, PA vs GB/PG | *RWDD2A, SPAG4, QPRT, PDLIM2, HS3ST2, REEP2, ITPKA, RABGEF1, UBXN10, HTRA1, ACHE, IMPDH1, RARRES2, STAC2, THY1, STC1, PRSS12, C11orf24, CEND1, EMP2* |
| H3K27ac, DA vs GB/PG | *ISG20, GLUD1, STC1, FOSL1, PDGFA* |

### Organization of chromatin into TADs contributes to expression of genes involved in gliomagenesis

We observed that distribution of signal fold changes between gliomas of different grades from all high-throughput experiments, including ATAC-seq, H3K4me3, H3K27ac ChIP-seqs, DNAse-seq, DNA bisulfite sequencing and RNA-seq followed the structure of TADs organization. We discovered that the fold changes of signals from all of those experiments computed for all three grade-differential-analyses were more homogenous within TADs, than among TADs (Table 2; Tables S1, S2). To validate the significance of the obtained correlation, we re-assigned these fold changes *in silico* to random TADs, maintaining the original numbers of genes per TAD, simulating the situation where positioning of genes in particular TADs would not matter. This led to a strong increase of the p values and drop of the H statistics (Table 2; Tables S1, S2). Such result confirms that the pattern of grade specific changes in chromatin activity and chromatin openness in gliomas is aligned with the segmentation of chromatin into topologically associating domains.

**Table 2.** Kruskal-Wallis test results for the original and permuted values of log2 (fold changes) between PA and GB/PG samples for different datasets.

| type of data | Kruskal-Wallis test H-statistic | p value | permuted 10x H-statistic (average) | permuted 10x p value (average) |
| --- | --- | --- | --- | --- |
| H3K27ac (promoters) | 6138 | $5.0 \times 10^{-249}$ | 2813 | 0.46 |
| ATAC-seq (promoters) | 5304 | $8.0 \times 10^{-157}$ | 2780 | 0.62 |
| DNA-me gene body | 5033 | $5.5 \times 10^{-133}$ | 2807 | 0.42 |
| H3K4me3 (promoters) | 4864 | $5.6 \times 10^{-114}$ | 2776 | 0.64 |
| RNA-seq | 4795 | $8.2 \times 10^{-110}$ | 2783 | 0.56 |
| DNase-seq (promoters) | 4433 | $6.7 \times 10^{-77}$ | 2784 | 0.61 |
| DNA-me promoters | 4193 | $6.1 \times 10^{-70}$ | 2711 | 0.54 |

Furthermore, we aimed to identify TADs which are particularly rich in DEGs and/or genes with DEMs in gliomas. We found a group of TADs significantly enriched in genes associated with glioma tumorigenesis (binomial distribution test, Benjamini-Hochberg corrected p < 0.05), to which we refer afterwards as ‘*glioma TADs*’. Those TADs were enriched for DEGs and genes with DEMs including H3K4me3, H3K27ac, DNA methylation, ATAC-seq, DNAse I-seq (Figure 2A). The far most enriched was TAD number 1101 localized at chr5: 140 660 415 - 141 580 433 (Figure 2B; Figures S2A-S2B).

**Figure 2.**
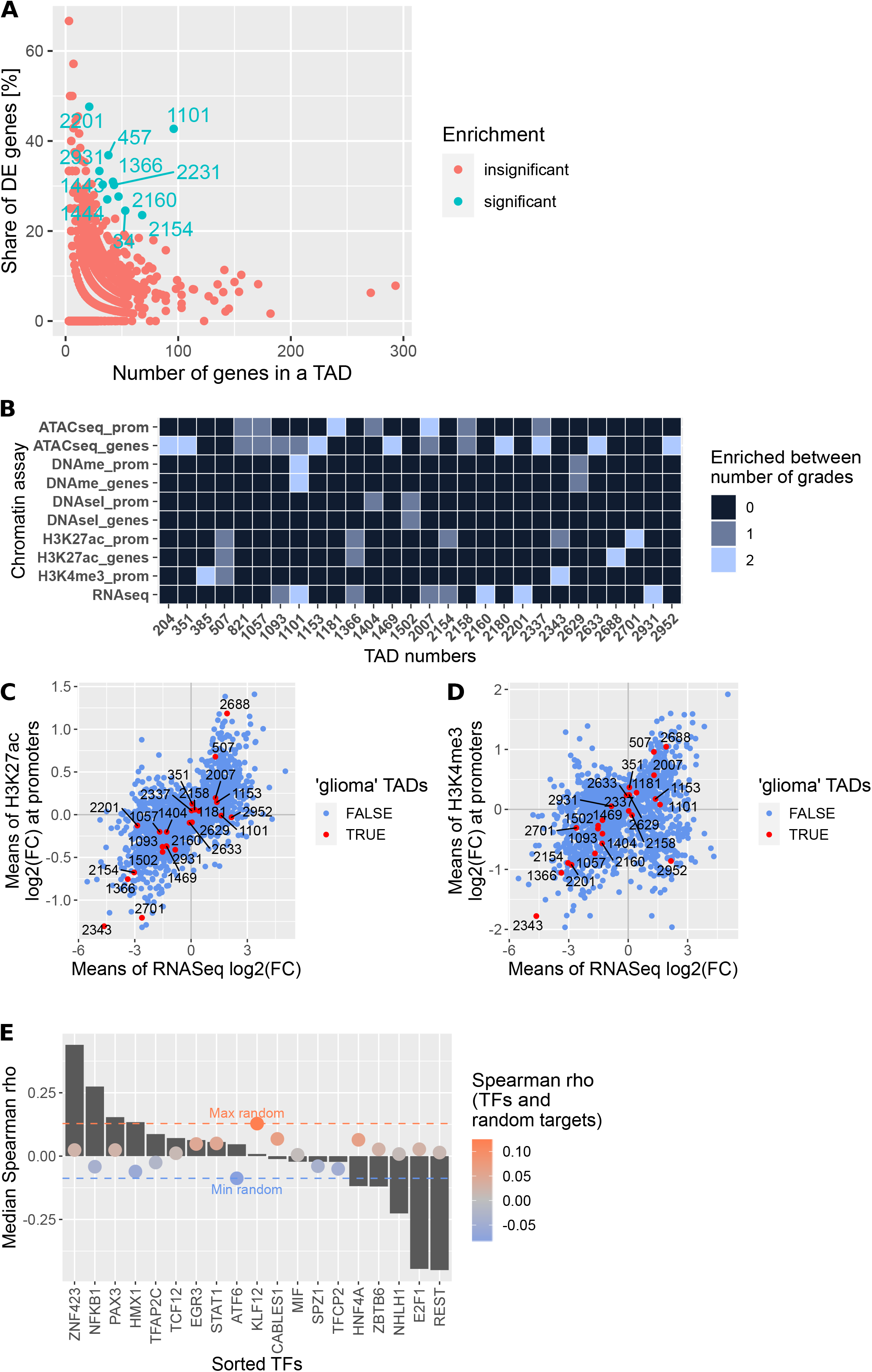
Discovering TADs enriched in genes differentially expressed and epigenetically modified in gliomas of different malignancies. (A) TADs enriched for DEGs or genes with DEMs deposited at their promoters, in the PA vs GB/PG comparison. (B) TADs with exceptionally high proportion of DEGs and genes carrying DEMs in all grades comparisons. Color scale depicts in how many of the three grade-comparisons a particular TAD was found to be enriched. (C) Dotplot showing means of DEGs expression and H3K27ac peak signals fold changes between PA and GB/PG samples. Each dot represents a mean for each TAD. Dots in red mark the most enriched TADs (‘*glioma TADs’,* binomial test Benjamini-Hochberg corrected p < 0.05). (D) As in Figure 2C for H3K4me3. (E) Spearman correlation between gene expression levels of enriched TFs and their target genes within TAD 2337. Grey bars depict median Spearman correlation between expression levels of genes encoding TFs and their target genes. Red and blue lines demarcate maximal and minimal correlation between expression levels of TF-coding genes and randomly selected, active genes. Colors of dots show median level of Spearman correlation between TFs and random target genes (red - positive correlation, blue - negative correlation).

Moreover, we observed that the fold changes of levels of H3K4me3 and H3K27ac deposition (measured as peaks’ heights) between different glioma grades correlated strongly with fold changes of DEGs expression at the TADs level (Figures 2C, 2D). For TADs containing DEGs between PA vs GB/PG, Spearman rho correlation coefficient for H3K27ac was 0.66 (p = 8.7 e-203). For the selected ‘*glioma TADs*’ the correlation was even stronger: Spearman rho = 0.86, (p = 2e-6). Similarly, for H3K4me3 and all TADs rho = 0.31, (p = 6.8e-37) while just for the *‘glioma TADs*’ rho = 0.75, (p = 4e-5). Similar results were obtained for the PA vs DA comparison (Figures S2C, S2D). This shows that the gene expression changes occurring between gliomas of different grades could be orchestrated by the TAD structure and are accompanied by the simultaneous changes in H3K4me3 and H3K27ac epigenetic marks deposition.

Genes found in the ‘*glioma TADs*’ play significant roles in cancerogenesis (Table 3; Figures S3A-S3D). Genes within the most enriched TAD 1101 encode cadherins which are involved in the cell adhesion process and Wnt signaling pathway. TAD boundaries and genes located within them are reported in the Table S3.

**Table 3.** TADs most enriched (enriched in at least 3 experiments for grade-differential comparisons, see Figure 2B) for ‘*glioma genes’* (‘*glioma TADs*’) and their functional annotations.

| TAD number | Biological Process GO terms | Enriched TFs common for the *glioma TADs*, (sign of correlation with target DEGs mRNA) |
| --- | --- | --- |
| 507<br>chr2:<br>221935281 -<br>222855282 | sphingosine metabolic process |  |
| 1101<br>chr5: 140660415<br>- 141580433 | synapse assembly,<br>cell-cell adhesion<br>(*PcdhgA* genes) | *E2F1* (negative), *REST* (negative), *PAX5* (negative), *NFE2L1* (positive), *NFKB1* (negative), *HMX1* (positive), *ZBTB6* (negative) |
| 1366<br>chr7: 27120381 - | cell differentiation<br>(*HoxA* genes) | *NFE2L1* (negative) |
| 28080381 |  |  |
| 2007<br>chr11: 62312528<br>- 62872528 | snRNA processing | *E2F1* (negative), *PAX5* (negative) |
| 2337<br>chr14: 23170791<br>- 24410794 | (cardiac) tissue<br>morphogenesis, G-protein<br>coupled receptor signaling | *E2F1* (negative), *REST* (negative), *NFKB1* (positive), *HMX1* (positive), *ZBTB6* (negative) |
| 2343<br>chr14: 28650794<br>- 29490794 | - |  |

To account for the potential effect of Copy Number Alterations (CNAs) on gene expression, we performed a CNA analysis using RNA-seq data. We found TAD 1366 to be amplified in 3 out of 12 analyzed GB samples (chr7 gain) and depleted in 2 GB samples and 3 out of 6 DA samples. The remaining TADs did not have amplified regions.

Further, we determined whether expression of DEGs present within the top six ‘*glioma TADs*’ (Table 3) could be regulated by the common transcription factors (TFs). For the sets of DEGs present within each of these TADs, we found enriched TFs using David tool (FDR<0.01). Then, for each pair of enriched TFs and their target DEGs we correlated their expression at the mRNA level. To validate the strength of these correlations, we also correlated expression levels of these TFs with randomly selected, active genes. Out of the original correlations we ultimately selected only those TFs whose median correlation values were higher than a maximal correlation (or lower than minimal in case of negative correlation) with random genes. In this way we obtained a list of 75 TFs that correlated strongly and significantly with their target genes (one example is shown for TAD 2337, Figure 2E) (Figures S4A-S4C; Table S4). The TFs with highest absolute values of correlation were: *ZNF423* (rho = 0.44), *REST* (rho = −0.45), *E2F1* (rho = −0.45) and *NHLH1* (rho = −0.23). We identified a group of TFs which putatively regulate gene expression in multiple TADs: E2F1, REST, PAX5, NFE2L1, NFKB1, HMX1 and ZBTB6 (Table 2). Positive regulation was found for known inducers of gene expression and negative for repressors. Additionally, the TF predictions were confirmed by an independent analysis with the BMO tool. We detected TF binding sites for E2F1, HMX1, PAX5, REST and ZBTB6 in the promoters of DEGs present within top six ‘*glioma TADs*’ (Table S5).

#### Glioblastoma-related TADs are enriched in DEGs with bivalent chromatin

Glioblastomas are described as frequently having marks of bivalent chromatin within gene promoters (Hall et al. 2018). In the studied patient cohort we found 54 DEGs (Table S6) with bivalent chromatin signals, simultaneously marked by H3K27me3 and H3K4me3 in GB samples (we did not obtain good quality H3K27me3 ChIP-seqs for PA and DA samples). Functions of these genes are associated with positive regulation of cell cycle, cell proliferation, cell adhesion and MAP kinase activity, among others (Figure S4D). Out of those genes, six were localized within three of the six most enriched ‘*glioma TADs*’: 2160 (*CDK4, TSFM, TSPAN31, B4GALNT1*), 2337 (*NFATC4*) and 2688 (*CACNA1G*). This shows that the TADs rich in genes involved in gliomagenesis are also enriched for the genes in bivalent chromatin state (p < 0.001, hypergeometric test).

### Regulation of DEGs transcription by long-range contacts with enhancers

#### Genes in contact with enhancers

We tested the hypothesis that in gliomas DEGs being in contact with enhancers will have elevated expression levels. Out of DEGs from PA vs GB/PG comparison, 41% (n=1716) got predicted long-range contacts with at least one enhancer. Those enhancers constituted 8.62% (n=3928) of all predicted enhancers in our samples. Some enhancers, however, did not have any predicted contacts and some had contacts with other loci than DEGs. DEGs with the predicted contacts with enhancers had significantly higher expression (by 11%) than DEGs for which such contacts were not assigned (Wilcoxon test, p=3.5-27).

#### Genes with multiple chromatin contacts

Recent findings showed that genes with multiple loops have usually higher expression than genes with single loops (Stevens et al. 2017, Johnston et al. 2019). Therefore, we identified genes having multiple contacts with enhancers localized on the same chromosomes (Figure 3A, Figures S5A, S5B). Out of DEGs from PA vs GB/PG comparison, 25% (n=1041) had predicted contacts with multiple enhancers (in PA vs DA 24% and in DA vs GB/PG 29%, respectively). Gene set enrichment analysis (GSEA) of those genes showed cell-cell adhesion via plasma membrane adhesion molecules (normalized enrichment score (NES): 1.9, q = 0.051) and stress-activated protein kinase signaling cascade (NES: 1.76, q = 0.065) as the most enriched sets. The most highly looped DEG was *PRDM16* which had 83 contacts with 24 enhancers (Figure 3A). Moreover, we found that one locus tends to be especially rich in long-range contacts with regions of undefined, possibly regulatory function. This locus contains *SAPCD1-AS1, MSH5, MSH5-SAPCD1, SAPCD1, LY6G6C, LY6G6D* and *ABHD16A* genes. Additionally, we confirmed that in glioma bulk tumor samples we could also observe a higher expression of DEGs having multiple (>=6) contacts with enhancers than just a few (1-5), (Wilcoxon test p = 1.1e^−17^, 2.1^−13^, 0.008 for PA vs GB/PG, PA vs DA and DA vs GB/PG, respectively, Figure 3B, Figures S5C, S5D).

**Figure 3.**
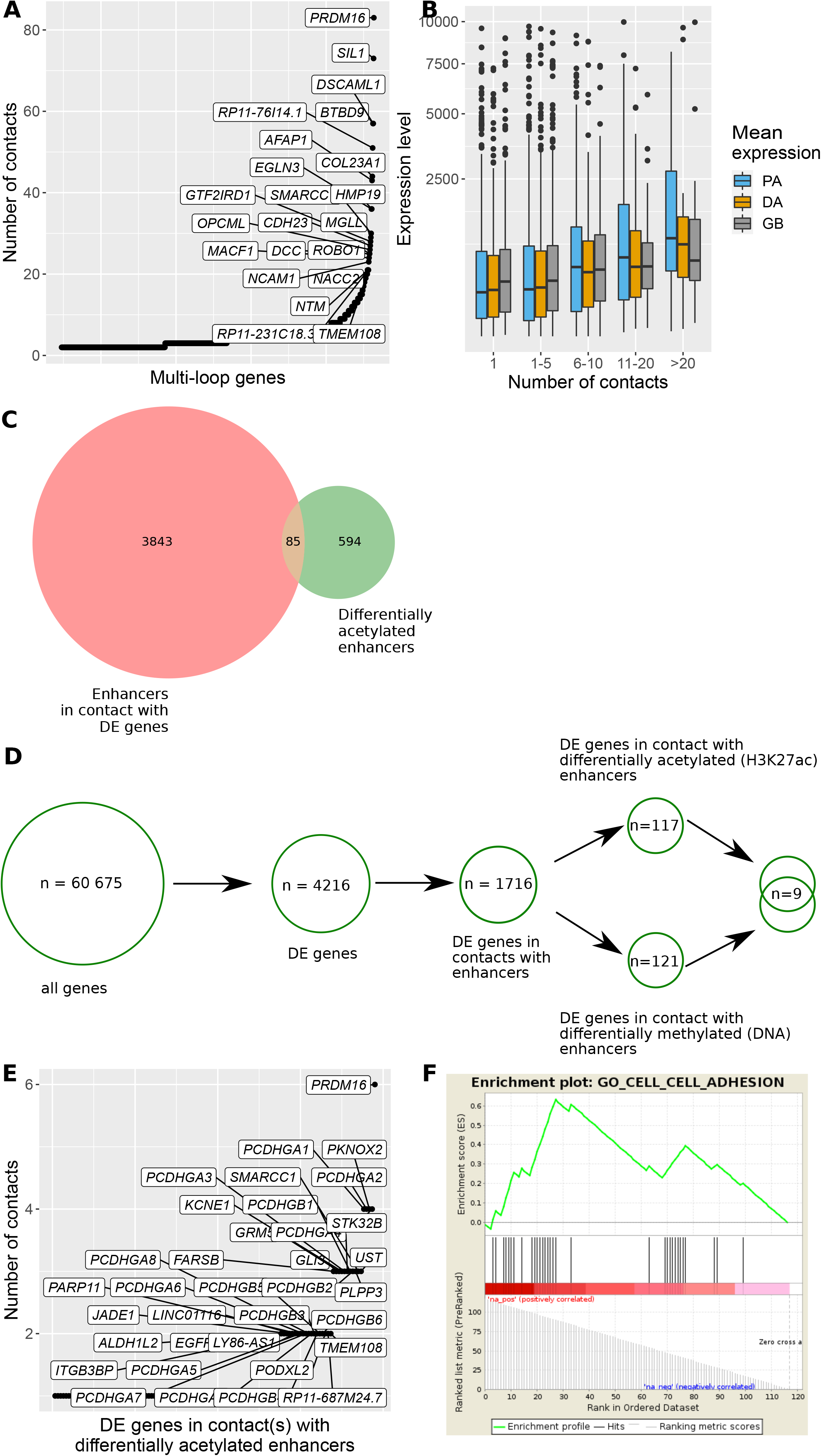
Prediction of gene regulatory networks enriched for multiple predicted contacts with enhancers. (A) DEGs from PA vs GB/PG comparison (n=1041) with predicted multiple contacts with enhancers. (B) Higher numbers of contacts with enhancers co-exist with higher expression of DEGs (PA vs GB/PG). (C) Enhancers contacting DEGs (PA vs GB/PG) intersect with the differentially acetylated (H3K27ac) (PA vs GB/PG) enhancers. (D) Workflow for obtaining DEGs potentially regulated by the differential activity of their contacting enhancers. The obtained enhancer activity might be due to differential H3K27ac and / or DNA methylation marks. Numbers are given for the comparison of PA vs GB/PG. (E) DEGs (PA vs GB/PG) contacting the differentially active enhancers (differential H3K27ac marks between PA vs GB/PG). (F) The most enriched gene set from GSEA performed on DEGs having at least 1 contact with differentially acetylated enhancers between PA vs GB/PG glioma samples.

To account for possible effect of CNAs on gene expression, we checked whether highly connected genes held CNAs. *PRDM16* did not have any CNAs, while the region around the *MSH5* gene was amplified in 3 out of 6 DA samples, in 7 out of 12 GB samples, but also in 4 out of 11 PA samples (among the PA samples there was also one deletion). The presence of CNAs is thus an unlikely reason for differential expression of these genes between PA and GB/PG. For the other highly looped genes, we found no effect of CNAs on their expression.

#### Enhancers with grade-specific activity

Following the observation that not all of the enhancers are active in all patients but they are rather patient-specific (Stępniak et al. 2020), we searched for the enhancers differentially active in glioma of different grades (Figure 3C; Figures S5E-S5F). We found 679 enhancers having significantly different levels of H3K27 acetylation between PA vs GB/PG samples (Wilcoxon test p <0.01). Out of them 85 (12.8% of the enhancers with contacts, Figure 3C; Figures S5E-S5F) were in contact with 117 DEGs (PA vs GB/PG) (Figure 3D). *PRDM16* had the highest number of contacts (n=6) with the differentially acetylated enhancer (chr1: 3078107 - 3079482) (Figure 3E). *PRDM16* gene encodes a methyltransferase which monomethylates histone H3K9. Moreover, there was a whole family of *PCDHGA* genes contacting a differentially acetylated enhancer (chr5: 141528260 - 141529748) (Figure 3E). The GSEA performed on all DEGs having at least 1 contact with differentially acetylated enhancers between PA and GB/PG showed significant enrichment of cell-cell adhesion genes (NES: 1.94, q = 0, Figure 3F).

#### Grade-specific enhancers may activate PROTOCADHERIN genes

Within the cell-cell adhesion set of genes there were found 14 *PCDHGA* genes that encode protocadherins - calcium-dependent cell-adhesion proteins. The cluster of those genes is located in one of the ‘*glioma TADs*’ - number 1101 (Table 3). *PCDHGA* genes showed overall high transcription in PA samples, lower in DA and lowest in GB and PG samples (Wilcoxon test p<0.0005, Figure 4A). The relatively low expression of those *PCDHGA* genes in PG, similarly to GB, shows their possible dysregulation related to cancer malignancy, not age. The level of H3K27 acetylation of the differentially acetylated enhancer contacting those genes was significantly higher in PA samples than in DA or GB/PG samples (Wilcoxon test, p = 0.07 PA vs DA, p = 0.003 PA vs GB/PG, Figure 4B). Interestingly, among the three enhancers contacting the group of *PCDHGA* genes, just one was differentially acetylated (Figure 4C), putatively affecting expression of the whole gene cluster. The median correlation between expression of *PCDHGA* genes with the acetylation of the enhancer (chr5: 141528260-141529748) across the samples was very high (Spearman rho = 0.73, p = 0.0002, Figure 4D). With use of BMO tool we found within this enhancer a presence of TF binding sites of multiple TFs: BACH1, FOSL1, IRX3, ZBTB33 (KAISO), NFATC3, SOX1, SOX21, YY1, ZFP42, ZNF418 and ZNF41 (Figure 4E). It is worth noticing that the highest mean positive correlation of expression of *PCDHGA* genes with the genes coding those TFs was observed for *SOX21* and *SOX1* (Spearman rho = 0.72 and 0.62, respectively, Figure 4E) and negative for *ZFP32* and *NFATC3* (Spearman rho = −0.57 and −0.42, respectively). We postulate that binding of those TFs within the chr5: 141528260-141529748 enhancer may influence expression of the *PCDHGA* gene cluster.

**Figure 4.**
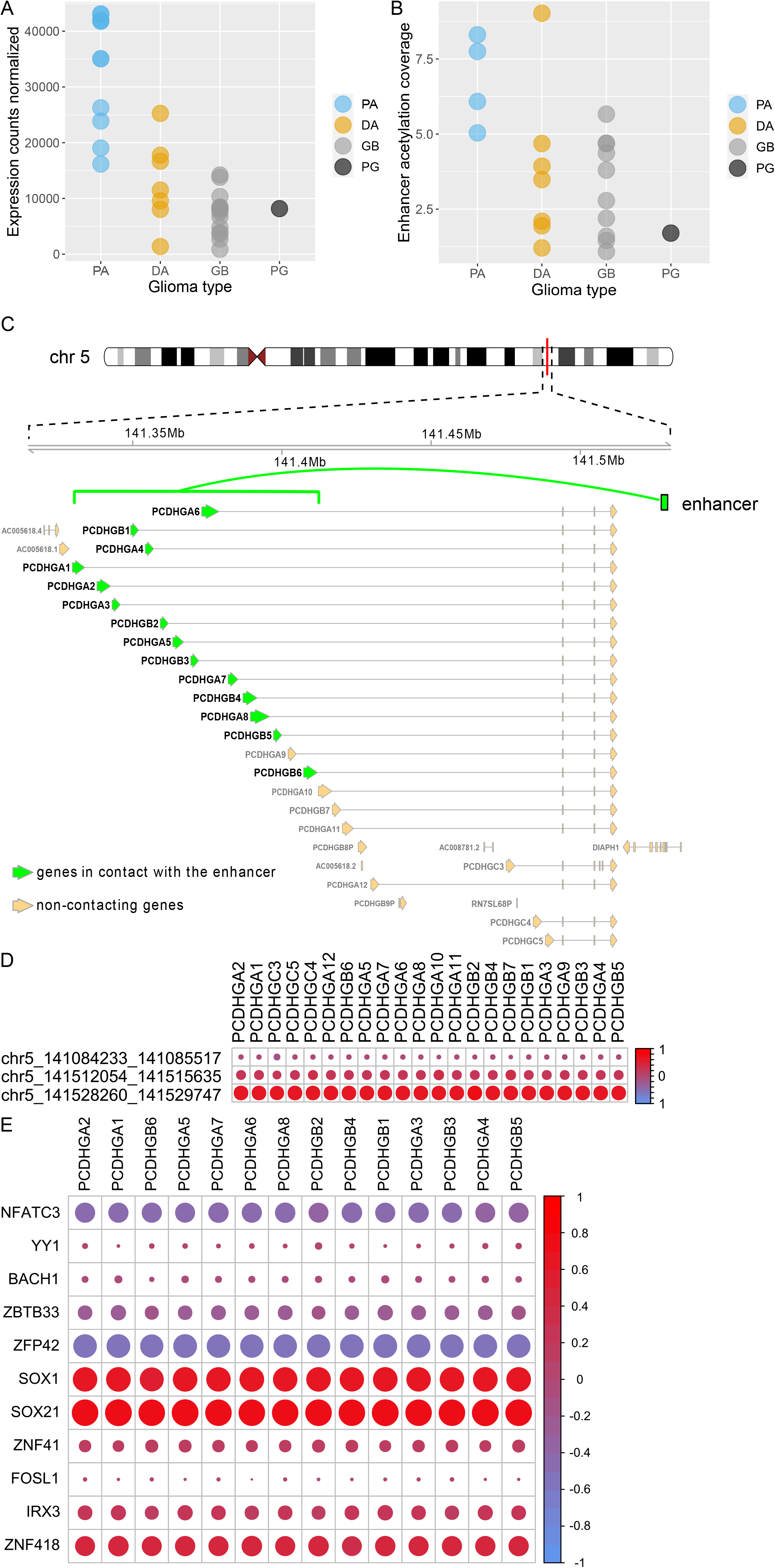
Regulation of *PCDHGA* gene cluster by H3K27 acetylation and/or DNA methylation of their contacting enhancers, as well as possible binding of transcription factors. (A) Expression of the *PCDHGA2* gene (as an example of *PCDHGA* genes which had similar expression levels) across glioma grades. (B) H3K27 acetylation levels at the enhancer (chr5: 141528260-141529748) contacting *PCDHGA2* gene across glioma grades. (C) Visualization of contacts between *PCDHGA* genes cluster and contacting enhancer at chr5: 141528260-141529748. (D) Correlations between expression levels of *PCDHGA* genes and H3K27ac levels of their contacting enhancers. (E) Correlations between expression levels of *PCDHGA* genes and TF-coding genes which binding sites were detected within the chr5: 141528260-141529748 enhancer. These TFs were identified using DAVID and BMO.

#### Enhancer-driven regulation deteriorates with tumor malignancy

All 117 genes contacting differentially acetylated enhancers show a strong correlation of enhancer acetylation with expression of target genes (Table 4). The observed correlation was most prominent for PA samples (Figure 5A, Table 4). To assess the significance of the obtained results, we performed a permutation test, in which DEGs were randomly paired with enhancers. The median correlation values dropped almost to zero (Table 4). Moreover, we observed that acetylation of enhancers contacting DEGs was generally higher in PA samples than in higher grade gliomas (DA/GB/PG) (Figure 5A). Interestingly, in higher grade gliomas enhancer acetylation was associated with the expression of the highly expressed genes, while in PA it was well pronounced across all RNA expression quantiles, increasing towards the upper quantile (Figure 5A). It shows that in DA, GB and PG samples high enhancer acetylation might enhance expression of already highly expressed genes. Overall, this result shows that a potential dysregulation of the gene-enhancers network might drive expression of genes important for glioma development.

**Table 4.** Correlation between expression levels of the 117 genes contacting differentially acetylated enhancers and acetylation peak heights of those enhancers.

|  | Spearman correlation rho (p) | Median Spearman rho for permuted pairs |
| --- | --- | --- |
| all grades together | 0.45 (0.04), sd = 0.34 |  |
| GI (PA) | 0.43 (1.1e<sup>-6</sup>) | -0.01, sd=0.09 |
| GII/III (DA) | 0.3 (0.001) | 0, sd = 0.1 |
| GIV (GB/PG) | 0.27 (0.003) | 0.02, sd = 0.1 |

**Figure 5.**
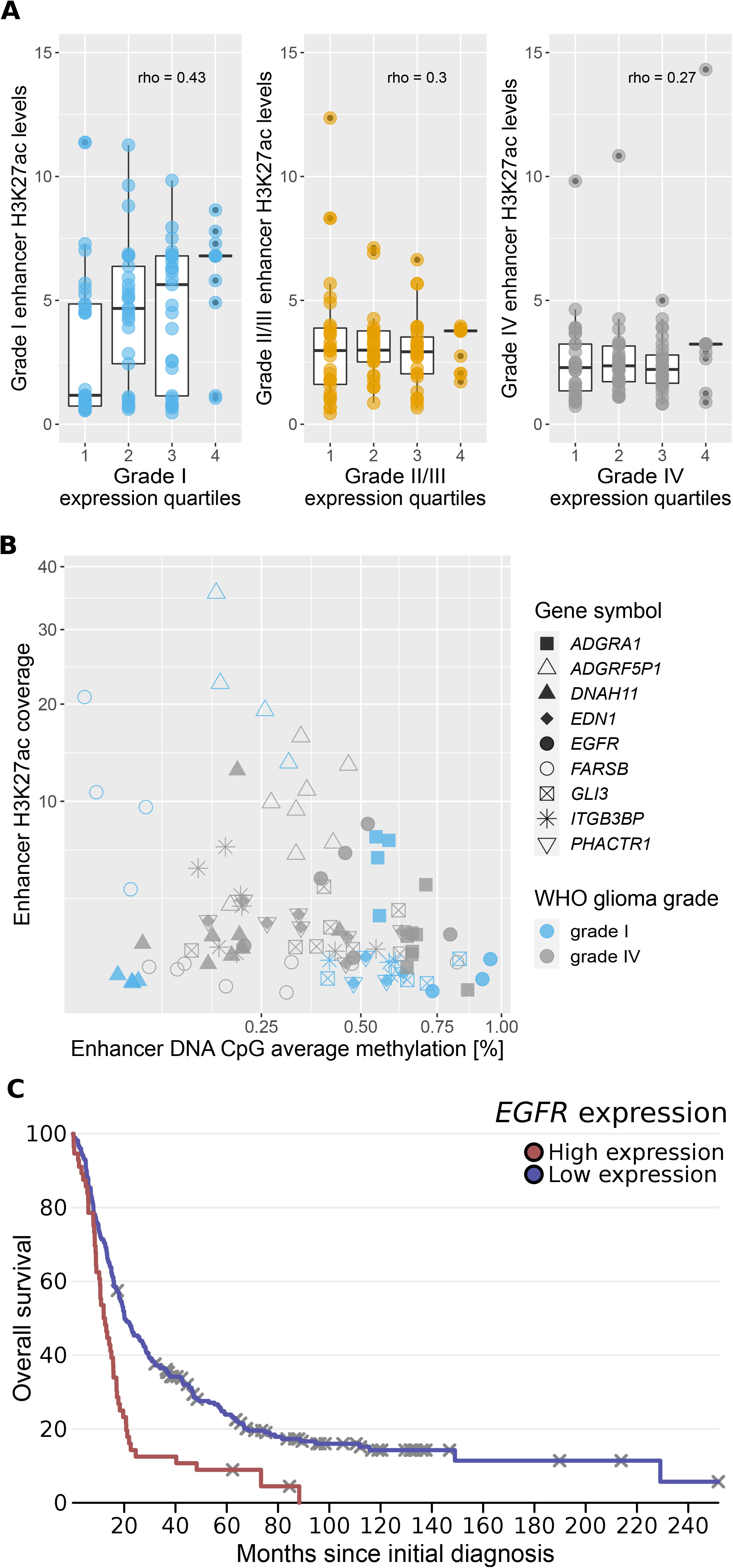
The effect of differential H3K27 acetylation and/or DNA methylation at enhancers on the expression of their target genes: *PROTOCADHERINS*, *EGFR* and other glioma-related genes. (A) Correlation between enhancers’ H3K27 acetylation and expression of DEGs (PA vs GB/PG) contacting them. Correlation decreases in higher grade gliomas. (B) Correlation between DNA methylation and H3K27ac signals at the enhancers being in contact with DEGs and being differentially methylated and acetylated (H3K27ac) between PA vs GB/PG samples. Each dot represents a gene in each sample - 9 genes in 12 samples for which there were simultaneously available good quality DNA methylation and H3K27ac data. (C) Survival analysis plot of the expression of *EGFR* gene showing its prognostic value for glioma patients survival, based on the Rembrandt dataset.

#### Activity of enhancers might be blocked by DNA methylation

Ultimately, for the DEGs having contacts with differentially acetylated enhancers (H3K27ac), we determined the role of DNA methylation of the contacting enhancers on gene expression. We found nine, out of 85, differentially acetylated enhancers having also differentially methylated DNA (Wilcoxon test, p<0.01, Figure 3D). The selected nine enhancers had predicted contacts with nine DEGs. For all those genes except the *DNAH11*, negative correlation between DNA methylation and H3K27 acetylation levels of enhancers was detected (median of Spearman rho calculated for all genes separately = −0.57, median p = 0.05, Figure 5B, for *DNAH11* Spearman rho = 0.79, p=0.03, Table S7). The enriched GO Biological Process terms for the nine genes were: heart and tongue development, cell surface receptor signaling pathway, dorsal/ventral pattern formation, positive regulation of nitric oxide biosynthetic process, positive regulation of MAP kinase activity and wound healing (David tool, p<0.05). One of the aforementioned genes was *EGFR*, which is a well-known cancer-related gene and its high expression predicts poorer survival in various cancers, including glioblastomas (Figure 5C). In our dataset *EGFR* was overexpressed in DA and GB samples compared to PA. At the same time, the enhancer (chr7: 54881593 - 54882643) related to the *EGFR* gene had significantly higher acetylation and lower DNA methylation in DA and GB/PG samples than in PA (p<0.01, Wilcoxon test; Spearman correlation of enhancer DNA methylation and H3K27 acetylation rho = −0.5, p = 0.17, n = 12 samples, Figures S5G, S5H). Half of the genes contacting differentially acetylated and differentially DNA-methylated enhancers, namely *GLI3*, *ITGB3BP*, *DNAH11*, *ADGRA1* and *ADGRF5P1* were found to be prognostic for patients survival (log-rank test p<0.0001, in the TCGA dataset for WHO GIII and GIV glioma patients; Table S7, Figures S6A-S6E). Higher expression of *GLI3*, *ITGB3BP* and *DNAH11* genes was associated with shorter survival, while higher expression of *ADGRA1* and *ADGRF5P1* – with longer survival. We excluded the single nucleotide polymorphisms (SNPs) as a potential reason for different levels of enhancer activation: none of the SNPs in a tumor DNA correlated with malignancy and had low population frequency (results not shown).

## Supplementary Tables

**Table S1.** Kruskal-Wallis test results for the original and permuted values of log2 (fold Changes) between PA (WHO GI) and DA (WHO GII/III) samples for different datasets.

| type of data | Kruskal-Wallis test H-statistic | p value | average H-statistic from 10 permutations | average p value from 10 permutations |
| --- | --- | --- | --- | --- |
| H3K27ac (promoters) | 5893 | $5.10 \text{ e}^{-208}$ | 2591 | 0.50 |
| ATAC-seq (promoters) | 5569 | $4.6 \text{ e}^{-173}$ | 2569 | 0.58 |
| DNA-me gene body | 4662 | $5.4 \text{ e}^{-89}$ | 2562 | 0.52 |
| H3K4me3 (promoters) | 5024 | $3.1 \text{ e}^{-119}$ | 2601 | 0.46 |
| RNA-seq | 4633 | $1.8 \text{ e}^{-88}$ | 2503 | 0.57 |
| DNase-seq (promoters) | 4022 | $7.1 \text{ e}^{-41}$ | 2600 | 0.46 |
| DNA-me promoters | 4701 | $2.1 \text{ e}^{-98}$ | 2506 | 0.37 |

**Table S2.** Kruskal-Wallis test results for the original and permuted values of log2 (fold Changes) between DA (WHO GII/III) and GB (WHO GIV) samples for different datasets.

| type of data | Kruskal-Wallis test H-statistic | p value | average H-statistic from 10 permutations | average p value from 10 permutations |
| --- | --- | --- | --- | --- |
| H3K27ac (promoters) | 6518 | $5.0 \text{ e}^{-280}$ | 2604 | 0.43 |
| ATAC-seq (promoters) | 6614 | $3.0 \text{ e}^{-292}$ | 2589 | 0.48 |
| DNA-me gene body | 4015 | $3.1 \text{ e}^{-41}$ | 2543 | 0.65 |
| H3K4me3 (promoters) | 5656 | $3.3 \text{ e}^{-182}$ | 2567 | 0.61 |
| RNA-seq | 4617 | $4.5 \text{ e}^{-85}$ | 2559 | 0.4 |
| DNase (promoters) | 6246 | $3.1 \text{ e}^{-248}$ | 2563 | 0.62 |
| DNA-me promoters | 3683 | $8.1 \text{ e}^{-26}$ | 2512 | 0.50 |

**Table S5.** Transcription factor binding sites detected within six top ‘*glioma TADs’* in the open chromatin regions using BMO tool on eight glioma samples of various grades.

| TADs numbers | TFBS |  |  |  |  |
| --- | --- | --- | --- | --- | --- |
|  | E2F1 | HMX1 | PAX5 | REST | ZBTB6 |
| 507 | 61 | 0 | 64 | 18 | 8 |
| 1101 | 16 | 0 | 30 | 0 | 20 |
| 1366 | 9 | 0 | 38 | 0 | 5 |
| 2007 | 146 | 0 | 137 | 43 | 51 |
| 2337 | 251 | 0 | 241 | 75 | 74 |
| 2343 | 0 | 1 | 2 | 0 | 0 |

---

**Figure S1 Identification of genes differentially expressed in gliomas of different malignancy grades - comparison of PA vs DA and DA vs GB/PG.**

(A) As in Figure 1A, for PA vs DA comparison.

(B) As in Figure 1A, for DA vs GB/PG comparison.

(C) As in Figure 1B, for PA vs DA comparison.

(D) As in Figure 1B, for DA vs GB/PG comparison.

(E) Enrichments for Biological Process GO terms for DEGs in PA vs DA comparison having high correlation of expression levels with H3K4me3 (Spearman rho > 0.7), and being prognostic for glioma patient survival (log-rank test < 0.001).

(F) As in Figure S1E, for H3K27ac.

(G) Enrichments for Biological Process GO terms for DEGs in DA vs GB/PG comparison having high correlation of expression levels with H3K4me3 (Spearman rho > 0.7), and being prognostic for glioma patients survival (log-rank test < 0.001).

(H) As in Figure S1G, for H3K27ac.

**Figure S2 Discovering TADs enriched for genes differentially expressed and epigenetically modified in gliomas of different malignancy grades - comparison of diffuse astrocytomas (DA) with lower and higher grades (PA, GB/PG)**

(A) TADs with exceptionally high proportion of DEGs and genes carrying DEMs in the PA vs DA comparison. Color scale depicts in how many of the three grade-comparisons a particular TAD was found to be enriched.

(B) As in Figure S2A, for DA vs GB/PG comparison.

(C) As in Figure 2C, for PA vs DA comparison.

(D) As in Figure 2C, for DA vs GB/PG comparison.

**Figure S3 Gene Ontology analysis of TADs with the high proportion of genes differentially active between different grades of gliomas**

(A) Enriched GO terms Biological Process categories for genes present within TAD 2007.

(B) As in Figure S3A, for TAD 507.

(C) As in Figure S3A, for TAD 1366.

(D) As in Figure S3A, for TAD 1101.

**Figure S4 Characterization of *‘glioma TADs*’ transcriptional regulation by transcription factors and functional annotation of bivalent chromatin genes**

(A) As in Figure 2E, for TAD 1101.

(B) As in Figure 2E, for TAD 1336.

(C) As in Figure 2E, for TAD 2007.

(D) Enriched GO terms Biological Process categories for genes having bivalent chromatin.

**Figure S5 Genes with multiple long-range contacts and enhancers targeting multiple genes - with the protocadherins example**

(A) DEGs from PA vs DA comparison with predicted multiple contacts with enhancers.

(B) DEGs from DA vs GB/PG comparison with predicted multiple contacts with enhancers.

(C) As in Figure 3B, for DEGs of PA vs DA.

(D) As in Figure 3B, for DEGs of DA vs GB/PG.

(E) As in Figure S3B (stara), for PA vs DA comparison.

(F) As in Figure S3B (stara), for DA vs GB/PG comparison.

(G) Dotplot illustrating EGFR expressions in patients of all glioma grades vs contacting enhancer H3K27 acetylation.

(H) Dotplot illustrating EGFR expressions in patients of all glioma grades vs contacting enhancer DNA methylation.

**Figure S6 Patients survival analysis for genes having differential enhancer H3K27 acetylation and DNA methylation between different glioma grades**

(A) Kaplan–Meier survival curve of *ADGRA1* high expression (red line) versus low expression (blue line) in WHO GIII and GIV patients, data from TCGA.

(B) As in Figure S6A, for *ADGRF5P1*.

(C) As in Figure S6A, for *DNAH11*.

(D) As in Figure S6A, for *GLI3*.

(E) As in Figure S6A, for *ITGB3BP*.

## DISCUSSION

Taking advantage of the unique datasets encompassing RNA-seq and different chromatin characteristics (H3K4me3, H3K27ac, H3K27me3, DNA methylation, ATAC-seq and DNAse-seq) datasets obtained from 33 gliomas of WHO grades I, II/III and IV, we studied the epigenetic mechanisms which might contribute to regulation of expression of genes involved in gliomagenesis. Our results showed that an interplay of chromatin organization and different epigenetic mechanisms provides a fine-tuned framework for regulation of gene expression which might lead to and/or sustain gliomagenesis.

We focused on the genes differentially expressed (DEGs) between benign and malignant gliomas and inspected the regulatory mechanisms underlying this differential expression. We demonstrated a stronger correlation between expression of the DEGs and H3K4me3 and H3K27ac histone marks known to denote active genes promoters (Calo and Wysocka, 2013, Engelen et al. 2015), than in comparable sets of non-glioma genes. Moreover, the patterns of glioma grade-related differences in epigenetic marks deposition at the promoters followed the 3-dimensional chromatin segmentation pattern into domains (TADs), as the variance of the observed epigenetic modifications or chromatin openness changes between glioma grades was higher between groups of genes belonging to different TADs than within TADs. This indicates that the within-TAD epigenetic changes occurring in a grade-related manner might be one of the factors regulating expression of gene clusters in glioma.

Another layer of regulation is provided by the long distance interactions such as ones occurring between genes and enhancers. It has been reported that in glioma stem cells genes forming multiple contact loops with enhancers, are more highly expressed than those not being in contacts (Johnston et al. 2019). We investigated whether such regulation takes place in glioma bulk tumor samples, and especially in differentially expressed genes, between gliomas of different malignancies. Therefore, we inspected which DEGs were predicted to have multiple long-range contacts with enhancers and we confirmed that genes having predicted multiple contacts had increased expression levels. We also identified a specific region on chromosome 6 (chr6: ~30 515 000 ~ 33 450 000) which was found to be very abundant in long-range contacts. This region did not have H3K27ac marks in our data and therefore, we could not definitely claim it is an enhancer, however its high connectivity suggests it might have a regulatory function. Interestingly, it is a gene-rich region containing genes which were predicted to be the most frequently contacted (40 - 97 predicted contacts) with another chromatin regions. The most contacted genes were localized within the Major Histocompatibility Complex (MHC) class III gene cluster, which is not well described yet. MHC class III genes are supposed to be involved in immunity and cell-signaling: genes for complement proteins, cytokines and heat shock proteins lie within the region (Gruen and Weissman, 2001; Milner and Campbell, 2001; Deakin et al. 2006). One of the most contacted genes, *MSH5*, encodes a MutS homolog, which is involved in homologous chromosome recombination (Rashambikai et al. 2013) and has been linked to DNA damage response and repair, neoplasia and immunity (Clark et al. 2013; Sekine et al. 2007). High expression of *MSH5* is associated with shorter survival of glioma patients (log rank test p = 1.08e-7, Rembrandt repository, Gusev et al. 2018) which is in agreement with our results. We found a higher *MSH5* expression in gliomas of WHO grades II/III and IV. The discovered group of genes within the MHC class III genes cluster might be extensively regulated by the long-range contacting regions. This group of genes requires further investigation and elucidation of their role in cancerogenesis.

Additionally, we found enhancers with H3K27ac signals significantly different in glioma of various grades - thus changing their activity along the malignancy grades. Correlation of enhancer H3K27 acetylation with the expression levels of their target genes ranged between 0.27 - 0.43 and was higher than in case of all active, non-DE genes in gliomas (Stępniak et al. 2020). Stępniak and coworkers have analyzed the same dataset and the correlation of expression of all active genes with H3K27ac of their contacting enhancers were rather low: 0.1 - 0.25. Altogether, these results suggest that there is a subset of genes revealed by this study which is strongly regulated by active enhancers and potentially contributes to gliomagenesis. The strength of the acetylation signal was very specific and related to particular malignancy grades.

The gene contacting the largest number of differentially acetylated enhancers (n=6) was *PRDM16*. This gene encodes a methyltransferase which monomethylates histone H3K9 (Pinheiro et al. 2012). *PRDM16* encodes a zinc finger protein which binds DNA and acts as a transcriptional regulator. *PRDM16* was over-expressed in leukemia samples (Nishikata et at. 2003) and is an important regulator of differentiation of myoblastic precursors into brown adipose cells (Seale et al. 2008). It can also act as a regulator of Transforming Growth Factor (TGF) β1 expression. TGFβ1 plays important roles in glioma proliferation, invasion and immunosuppression and is aberrantly expressed in gliomas (Kaminska, Cyranowski 2020). We demonstrate higher expression of *PRDM16* in high grade gliomas. Differential H3K27 acetylation, across grades, of the enhancers contacting the *PRDM16* gene can be thus the factor affecting the *PRDM16* expression and may contribute to gliomagenesis.

DNA methylation at CpG sites usually leads to transcription silencing and shows a negative correlation with enrichment of histone marks at enhancers (Calo and Wysocka, 2013). We searched for glioma-grade specific pairs of enhancers-target genes, in which activities of enhancers activity could be impaired by DNA methylation, resulting in weaker activation of their target genes transcription. Indeed, we found an inverse correlation of CpG methylation and H3K27 acetylation at enhancers contacting DEGs. Interestingly, among the genes contacting differentially acetylated (H3K27ac) and differentially methylated (DNA) enhancers, we found the *EGFR* gene. It encodes an epidermal growth factor receptor which controls cell proliferation, is implicated in cancerogenesis (Verhaak et al. 2010, Yarden et al. 2001, Nicholson et al. 2001) and its high expression negatively correlates with glioma patient survival. *EGFR* expression was found to be elevated in gliomas (Verhaak et al. 2010, Venneti and Thompson, 2012) and in our dataset *EGFR* was upregulated in high grade gliomas, accordingly. Apart from the differential H3K27 acetylation of the enhancers, we also uncovered that the poorly acetylated enhancers are characterized by high DNA methylation, suggesting that these two molecular mechanisms might be both involved in the regulation of *EGFR* expression.

In our study, we also identified cues for epigenetic regulation of a significant group of *PCDHG* genes in gliomas. *PCDHGA* genes encode protocadherins involved in cell–cell adhesion, boundary formation and maintenance as well as morphogenesis in the developing brain (Halbleib and Nelson, 2006). It has been shown that certain clusters of *protocadherins*: alfa (*PCDHA*) and gamma (*PCDHG*) are regulated in the stochastic manner in the developing human neurons (Almenar-Queralt et al. 2019). The stochastic *protocadherin* selection occurs at the level of neuronal progenitors and *protocadherin* expression is reduced or silenced in adult brain neurons. Chromatin activation (observed as H3K4me3 mark depositions) of the *protocadherin* promoters controls the stochastic regulation of their expression (Almenar-Queralt et al. 2019). Moreover, it has been shown that epigenetic dysregulation of *protocadherin* gene clusters might be linked with psychiatric and neuronal disorders (El Hajj et al. 2017). Protocadherins play a significant role during cancerogenesis. Decrease or loss of protocadherin expression has been correlated with poor prognosis in breast cancer patients (Berx and Van Roy, 2001). DNA methylation at the *protocadherins* promoters was observed to decrease their expression in many types of cancers (Sui et al. 2012). Such a decrease of expression has been reported also in high grade gliomas (Burgess et al. 2008). Here, we demonstrated the highest expression of the *PCDHG* genes in PA (WHO grade I) samples, moderate in DA (WHO grade II/III) and the lowest in GB and PG (WHO grade IV). Those genes exhibited high correlation of transcriptional activity with H3K4me3 and H3K27ac deposition at their promoters, which is in concordance with previously reported results (Almenar-Queralt et al. 2019). Moreover, we predicted contacts of those genes with enhancers located up to 2 Mb away from the promoters, including a differentially acetylated (H3K27ac) one. H3K27ac deposition at the non-promoter regions is usually associated with active enhancers (Creyghton et al. 2010). The highest H3K27 acetylation at enhancers contacting the *PCDHG* genes was found in PA and correlated well with their increased expression. Thus, we identified a new, potential mechanism in which the expression of the *PCDHG* gene cluster is regulated by acetylation of the contacting enhancer. Long-range regulatory elements were earlier reported but only for *protocadherin-alpha* gene cluster (Ribich et al. 2006). Here we presented the enhancer of *protocadherin-gamma* cluster which to our best knowledge is the first reported case. Additionally, the *PCDHGA* gene cluster was localized within the most enriched TADs in glioma DEGs.

Further, we predicted which transcription factors potentially regulate the cluster of the *PCDHG* genes by analyzing occurrence of transcription factor binding motifs within their promoters. Among the enriched TFs and most highly negatively correlated with the *PCDHG* cluster genes expression were E2F1 and REST, which are known as transcriptional repressors (Martínez-Balbás et al. 2000, Bruce et al. 2004). E2F1 expression was previously reported to be upregulated in different human cancers (Polager and Ginsberg, 2009) and we corroborated its increased expression in glioma of higher grades. Moreover, we found a positive correlation of *PCDHG* gene cluster expression with two other transcription factors LMO2 and NR2F1 which can act as transcription activators or repressors depending on their interacting partners (Wu et al. 2018, Shibata et al. 2003). Intriguingly, the positive correlation was also found with KLF12 which was reported to act as a repressor (Black et al. 2001).

Overall, the results shown here demonstrate the existence of epigenetic differences associated with chromatin organization driving differential gene expression in gliomas of different malignancy. We demonstrated that integration of whole genome, high-throughput, epigenetic data with Hi-C data and transcriptomic profiles can reveal new regulatory networks in low and high grade gliomas.

## MATERIALS and METHODS

### Samples and wet lab experiments

In our study we analyzed data published by Stępniak et al. 2020 on glioma samples which included gliomas assigned to different malignancy groups: pilocytic astrocytomas (PA, WHO grade I, n = 11), diffuse astrocytomas (DA, WHO grade II or III, n = 7), glioblastomas (GB, WHO grade IV, n = 14) and pediatric glioblastoma (PG, WHO grade IV, n = 1). PA and PG samples were derived from pediatric patients.

We studied the chromatin openness state (using ATAC-seq, DNAse-seq), deposition of epigenetic marks characteristic for open (H3K4me3, H3K27ac) or closed chromatin (H3K27me3) using ChIP-seq assays and DNA methylation (whole genome bisulfite sequencing) in the glioma samples as well as gene expression (RNA-sequencing) in freshly isolated gliomas of various grades. The detailed description of the samples and laboratory procedures is available in the publication by Stępniak et al. (2020).

### Genes with differential expression / epigenetics marks deposition

Differentially expressed genes (DEGs) were determined using DeSeq2 (Love et al. 2014), with false discovery rate (FDR) correction for multiple testing and significance level threshold FDR<0.01. Differential analysis with DeSeq2 was performed for three pairwise comparisons: i) WHO GI pilocytic astrocytomas (PA) vs WHO GII/GIII diffuse astrocytomas (DA) (PA vs DA), ii) PA vs WHO GIV glioblastomas (GB) and pediatric glioblastoma (PG) (PA vs GB/PG) and iii) DA vs GB/PG. Similarly as DEGs, genes with differential epigenetic marks (DEMs) were identified using DeSeq2 and FDR<0.01. Using different assays, we identified genes having grade-differential signals at their promoters or gene bodies. For promoters, defined as TSS+/− 2 kb, we used data from H3K4me3 and H3K27ac, ATAC-seq, DNA methylation and DNAse-seq assays. For gene bodies, we used DNA methylation assays. For further steps of our analytical pipeline, we selected DEGs and genes with assigned DEMs.

Correlation of gene expression and epigenetic marks within the promoters of DEGs or genes with DEMs was calculated with Spearman correlation, across patients. To compare correlation strength between gene expression and epigenetic marks among DEGs and randomly chosen active genes (>10 reads on average in all samples), we selected sets of genes having on average the same expression levels as in the original set of DEGs. For all the genes throughout this study we used their canonical TSS.

### Permutation testing for the significance of the overlap between genes with differential expression and differential deposition of the epigenetic marks

To verify the significance of the overlap of DEGs and genes with DEMs, we performed permutation testing (Figure 1A). Out of a pool of active genes, defined as those having the mean count across samples > 10, we randomly selected genes matching the number of: (1) obtained DEGs and (2) obtained genes with DEMs, and the overlap between them was calculated. Then, a random selection out of all, active genes of two sets of genes matching numbers of DEGs and genes with DEMs was repeated 100 times. In the next step, we calculated how many times the overlap of the randomly selected DEGs and genes with DEMs was equal or greater than the number of the originally overlapping genes. Based on the permutation test we computed the probability of obtaining the original number or greater of overlapping genes by chance.

### Searching for correlation between H3K4me3 and H3K27ac marks and gene expression

For DEGs, correlation between expression levels and H3K4me3 and H3K27ac levels across samples was calculated (Figure 1B). We compared the obtained correlation for DEGs with their epigenetic marks to randomly selected non-DEGs with their matching epigenetic marks. The randomly selected non-DEGs were sampled from active genes (average expression levels above 10) under the condition that their mean expression level was at least the same as that of the DEGs, to avoid possible bias resulting in stronger correlation due to stronger expression. Correlation was calculated across all samples of our dataset, separately for each of the pairwise comparisons: PA vs DA, PA vs GB/PG, DA vs GB/PG. Random selection of active genes matching numbers of each of the two groups was repeated 10 times for each epigenetic mark and grade pairwise comparisons.

We calculated a share of DEGs highly correlating with epigenetic marks among genes having a prognostic property for survival of glioma patients according to proteinatlas.org (Table S8_“proteinatlas.tsv”). The overlap between the highly correlating DEGs with protein atlas prognostic genes was divided by the total number of DEGs. Genes were estimated as having a prognostic value if they passed the log-rank test with p-value below 0.001 using dataset (Table S8_“proteinatlas.tsv”) from proteinatlas.org (Uhlen et al, 2015).

### Distribution of chromatin states across topologically associating domains

We investigated whether chromatin segmentation into TADs specifically affects DEGs and genes with DEMs involved in gliomagenesis (‘*glioma genes*’) Next, we searched for TADs particularly rich in ‘*glioma genes’*. This was done using a binomial distribution test with Benjamini-Hochberg corrected p < 0.05 set as significance threshold. We used Hi-C data generated on the mid-gestation developing human cerebral cortex with the segmentation of chromatin into 3165 TADs (Won et al. 2016).

Genomic coordinates of all genes (region from gene start to gene end defined as gene body) were acquired from human reference genome genome-build-accession NCBI:GCA_000001405.20 (hg38). Promoter regions were defined as +/− 2kB from TSS. For so defined promoters, data on deposition of H3K4me3, H3K27ac, open chromatin (ATAC-seq and DNAse-seq) and DNA methylation were assigned. Genes and their epigenetic marks were assigned into TADs genomic coordinates using Hi-C data (Won et al. 2016). To test co-regulation of different epigenetic marks as well as gene expression within the TADs, we performed the Kruskal-Wallis test. As the input for Kruskal-Wallis test log2 fold changes were calculated using DeSeq2 for all genes (including non-DEGs) or epigenetic marks assigned to genes. Differences using DeSeq2 were computed between: PA and DA, PA and GB/PG, DA and GB/PG. In the Kruskal-Wallis test calculated log2 fold changes were used as values and TAD borders as groupings. We used the fold changes of counts in promoter regions (TSS +/− 2kb) in case of H3K4me3 and H3K27ac, DNA methylation, ATAC-seq and DNAse-seq, as well as counts from the gene bodies for DNA methylation, ATAC-seq and DNAse-seq. To verify the significance of the Kruskal-Wallis test performed on the real data, we applied permutation tests. Permutation test was performed on data from each of the high-throughput measurements and within each of the group pairwise comparisons. In the permutation tests we shuffled the genes expression values / epigenetic marks values within the TADs, maintaining the original numbers of genes in each TAD. We performed the permutations 10 times for each type of data (e.g. ATAC-seq-gene bodies) and grade comparison. For all the tests we calculated the Kruskal-Wallis H statistics and p values. The original H statistics and p values were always at the top (or bottom, respectively) of the ranked, permuted values.

### Identification of TADs enriched for DEGs/DEMs

To identify TADs enriched in DEGs/DEMs, first we assigned them to TADs using custom script in R. For each TAD we calculated the probability of obtaining the observed number of DEGs/DEMs or higher, using a pbinom function from R ‘stats’ package, with input values being: q = (number of DEG/DEMs −1); size = number of genes in the TAD; prob = share of DEG/DEMs among all genes, lower.tail = F. Gene-poor TADs containing less than any three genes were excluded from the binomial distribution testing. The p-values were corrected using a Benjamini-Hochberg procedure (Benjamini and Hochberg, 1995). The whole procedure was repeated for all types of high-throughput experiments measurements and grade-differential pairwise comparison. The TAD borders were derived from Won et al. 2016.

We aimed to find a relationship between differential gene expression and epigenetic marks deposition in TADs. To find possible association between co-occurring changes in gene expression levels and H3K27ac/H3K4me3 depositions, we calculated for each TAD the mean gene expression fold changes of the genes present within a given TAD as well as the mean H3K27ac/H3K4me3 deposition fold changes. Ultimately, we correlated fold changes of gene expression levels within TADs between 3 pairwise comparisons of grades with their matching epigenetic marks fold changes obtained for the same TADs. Functions of genes present in the most enriched TADs were assigned using tool Enrichr (Chen et al. 2013).

### Identification of TFs potentially regulating DEGs present within the ‘*glioma TADs*’

In the above presented analyses we identified TADs enriched in DEGs/DEMs and called them ‘*glioma TADs*’. To determine whether expression of DEGs localized within the most enriched ‘*glioma TADs*’ may be regulated by specific TFs, we first searched for the enriched TFs for DEGs present within those TADs using David tool and FDR<0.01 cut-off. Then, for the pairs of enriched TFs and their DE target genes we calculated correlation of their expression. To validate the strength of correlation, we computed correlation of the same TFs with randomly selected, active genes. Then, we considered only those TFs which had median correlations values higher than maximal (or lower than minimal in case of negative correlation) correlation with random genes.

Additionally, prediction of TFs binding sites from DAVID tool was verified by an additional analysis of ATAC-seq data (Stępniak et al. 2020) employing BMO tool - a method to predict TF binding sites without using footprints (Albanus et al. 2019). First, low-quality reads and PCR duplicates were removed and only properly paired and uniquely mapped reads were retained for downstream analysis. Next, ATAC-seq peaks were called using MACS2 (Gaspar, 2018) using “broad” and “nomodel --shift −100 --extsize 200” flags. Resulting peaks were then intersected with human ENCODE blacklist regions (hg38) to discard the peaks within artifact regions. Concurrently, FIMO tool (Grant and Bailey, 2011) was used to predict occurrence of known motifs across the human genome (fasta file). Altogether, BMO pipeline was run and only the significant binding motifs instances, based on adjusted Benjamini-Yekutieli test (-log10 adjusted p-value < 0.05), were selected.

For the 315 genes encompassed within ‘*glioma TADs*’ (Table 2) we specified which TFBS are present within their promoters using the BMO tool. Gene promoters were defined as a sequence of 2 kb +/− from TSS. Detection of TFBS with BMO was performed using ATAC-seq from eight tumor samples (PA n=4, DA n=2, GB n=2). TFBS obtained with BMO allowed us to select TFs which could potentially regulate expression of the selected genes. Finally, TFBS were also detected within the enhancer region (chr5:141528260-141529748) associated with *PCDHGA* genes cluster with use of BMO.

### TADs with genes with bivalent chromatin

To investigate the presence of bivalent chromatin in the TADs defined as ‘*glioma TADs*’ we searched for genes characterized by H3K4me3 and H3K27me3 marks at their promoters. The H3K27me3 ChIP-seq data at the acceptable quality at our disposal were only from GB samples. We searched for the genes with the highest H3K4me3 and H3K27me3 ChIP-seq peaks (on average across GB samples). We defined bivalent chromatin as being simultaneously marked by active H3K4me3 and repressive H3K27me3 signals in the gene promoters. We sorted genes according to their H3K4me3 and H3K27me3 signals and selected 1000 with the highest signals in each group (>27.000 counts per sample, on average for H3K4me3 peaks; (>38.000 counts per sample, on average for H3K27me3). We assigned the genes to TADs and used hypergeometric distribution test to calculate the probability of the obtained overlap among the ‘*glioma genes’*.

### Long-range intra-chromosomal contacts of DEGs and enhancers

Following the finding of Johnston et al. 2019 that genes active in glioma stem cells with multiple loops have usually higher expression levels, we tested whether a similar pattern could be found in bulk tumor samples. Identification of enhancers, long-range contacts between genes and enhancers, as well as DNA methylation levels was previously described (Stępniak et al. 2020). In brief, active enhancers were determined by the presence of H3K27ac peaks in non-promoter regions, (with promoters defined as TSS +/− 2kb). We selected contacts between DEGs and enhancers within the 2 Mb range based on chromatin contact maps generated from the Hi-C data from developing human brains (Won et al. 2016). The differentially acetylated enhancers and differentially methylated CpGs within enhancers were defined by Mann-Whitney-Wilcoxon test p-value cutoff < 0.01. Enrichment scores for genes having multiple contacts with enhancers on the same chromosomes and for DEGs having at least 1 contact with differentially acetylated enhancers between PA and GB/PG were calculated using gene set enrichment analysis (GSEA, Subramanian et al. 2005). Other GO terms enrichments were done with DAVID tool (Huang et al. 2009). GO terms enrichments were visualized with the enrichR R package (Chen et al. 2013). Calculations of correlation, statistical tests and permutation testing was done using R (version 3.4). Genes for GSEA were ranked according to their number of loops.

### Correlations of DEGs expressions and DNA methylation or histone acetylation of grade-specific enhancers

We have calculated correlation of DEGs expression and their enhancers acetylation (H3K27ac) and DNA methylation. Enhancer assignment to the target gene was described in the previous paragraph. Permutation test was applied to test the significance of the correlation between DEGs expression and H3K27ac marks deposited at the contacting enhancers. For each grade separately we randomly paired DEGs expression levels and H3K27 acetylation at enhancers and calculated correlation. The procedure was repeated 100 times and means were calculated. For correlation with DNA methylation for each enhancer we used the average of CpG methylation levels localized within that enhancer.

### Patients’ survival analysis

Prognostic value of the *EGFR* gene expression for the patients’ survival was assessed using the Rembrandt repository - a large collection of genomic data from brain cancer samples (Gusev et al. 2018) and for all the other genes important for the study using the TCGA dataset (TCGA Research Network: https://www.cancer.gov/tcga).

### Copy number alterations analysis

In order to identify and visualize copy number variations (CNVs) in gliomas, we employed CaSpER (Harmanci et al. 2020), a signal processing tool that uses RNA-seq information to detect focal and large-scale genetic variations. B-allele frequencies were generated from RNA-seq pileup data to decipher allelic imbalances in PA, DA and GB samples. Concurrently, NGS reads were summarized at the gene level using featureCounts, in paired-end and reversely stranded modes (Kent et al. 2002) and resulting raw data were normalized to TPMs. After that, CNVs were called following the CaSpER tool author's guidelines.

### DE genes cross-validation with TCGA dataset

Identification of DE genes in gliomas of WHO grade II/III vs WHO grade IV in TCGA dataset consisted of calculation of the Wilcoxon test for each gene and applying Bonferroni correction. For DE genes from TCGA and our dataset we used a corrected p cutoff < 0.01.

Scripts written in R or python allowing to reproduce the results are deposited at https://github.com/ilona-grabowicz/epigenetics-in-glioma

## Acknowledgements

The authors thank Jacek Koronacki for his comments on the manuscript and Michał Dramiński for his contributions to optimizing computations. IEG, BWilczyński, BK, MJD have been supported by Polish National Science Centre grant [DEC-2015/16/W/NZ2/00314]. BWojtaś, AJR, BK were supported by the Foundation for Polish Science TEAM-TECH Core Facility project “NGS platform for comprehensive diagnostics and personalized therapy in neuro-oncology”.

## Authors contributions

IEG, BWilczyński, BK, BWojtaś and MJD conceived the study. IEG designed the study, performed majority of the bioinformatics and statistical analyses, interpreted and visualized results and wrote the original draft; IEG, BWojtaś and MJD performed additional bioinformatic analyses, review and editing; AJR performed CNAs and BMO analyses. All authors reviewed and edited the manuscript.

